# Hidden Markov Model-Based Prokaryotic Genome Space Mining Reveals the Widespread Pervasiveness of Complex I and Its Potential Evolutionary Scheme

**DOI:** 10.1101/2025.03.13.643058

**Authors:** Akshay Shirsath, Snehal V. Khairnar, Abhirath Anand, Divya M. Prabhakaran, Amitesh Anand

## Abstract

Most cellular reactions are interdependent; however, a subset of reactions often associates more closely to form a defined reaction pathway. An extreme arrangement of interdependent reactions occurs when the cognate proteins physically associate to constitute a complex. Respiratory complex I is one of the largest membrane resident protein assemblies. Besides being the hallmark of bioenergetics, this enzyme complex is critical for redox homeostasis and transport. The evolutionary scheme for the development of this enzyme complex is poorly understood due to associated challenges like complications in delineating close homologs and diverse subunit ancestry. We used custom Hidden Markov Model profiles to examine the available prokaryotic genome space to trace the distribution pattern of fourteen core Nuo subunits of Complex I. We report: (a) a sensitive HMMER-based workflow for comprehensively annotating and analyzing the Nuo subunits, which can be adapted to multiple such analyses; (b) the first curated species-level distribution of Nuo subunits; (c) multiple variants of Complex I across ∼11,000 species with 51.2% species having complete complex; (d) presence of Complex I variants on plasmids which potentially facilitated the evolutionary distribution; (e) extension of our workflow for examining distribution of mitochondrial Complex I accessory subunits among prokaryotes highlighting their evolutionary roots. We have also developed a web application to facilitate the convenient dissemination of our compiled resources. The knowledge of bioenergetic repertoire is critical in the successful targeting of energy metabolism for antimicrobial development.

## Introduction

NADH-quinone oxidoreductase (Nuo), commonly known as Complex I, is one of the most sophisticated assemblies of proteins wherein various subunits come together to form a relay system to transfer electrons from NADH to respiratory quinones. This multi-step quantum mechanical electron tunneling is achieved by a non-covalently bound flavin mononucleotide and multiple iron-sulfur clusters spread across various subunits of the N- (NADH dehydrogenase module) and Q- (quinone hydrogenase module) modules of Complex I ^1,2^. The placement of multiple redox centers across a distance of approximately 95 Å allows the hopping of electrons between NADH and quinones at biologically appropriate electron transfer rates. The reduction of respiratory quinones occurs at the Q-module. The peripheral N- and Q-modules interact with the membrane-embedded P-module (proton translocation module). The P-module comprises four proton channels that pump protons to the periplasmic space by coupling reduction potential favored electron flow in N- and Q-modules.

Complex I is a highly conserved enzyme complex in organisms ranging from bacteria to humans ^3^. This enzyme complex is a key component of the electron transport system (ETS), playing a critical role in cellular respiration. Remarkably, Complex I is thought to have evolved primarily to mitigate the pH stress of ancient fermentative metabolism ^4^. Mutations in this complex have recently been shown to alter intracellular acidity, leading to antibiotic persistent metabolic state ^5^. Such physiological implications suggest a deep and extensive evolutionary history of Complex I. While mitochondrial Complex I has acquired many accessory subunits, the fourteen core Nuo subunits are conserved across lifeforms ^6^. These core Nuo subunits share common ancestry with various protein families and are homologs of enzymes performing similar standalone catalytic functions (Figure 1A) ^7,8^. The soluble [Ni-Fe] hydrogenases are phylogenetically related to the N-module and the connecting Q-module subunits. The P module’s subunits show homology with the Na^+^/H^+^ antiporter MrpA and MrpC ^9^. Therefore, while intuitively, the emergence of Complex I appears to be a synergistic association of independent proteins, the actual evolutionary course has remained unclear.

**Figure 1:**
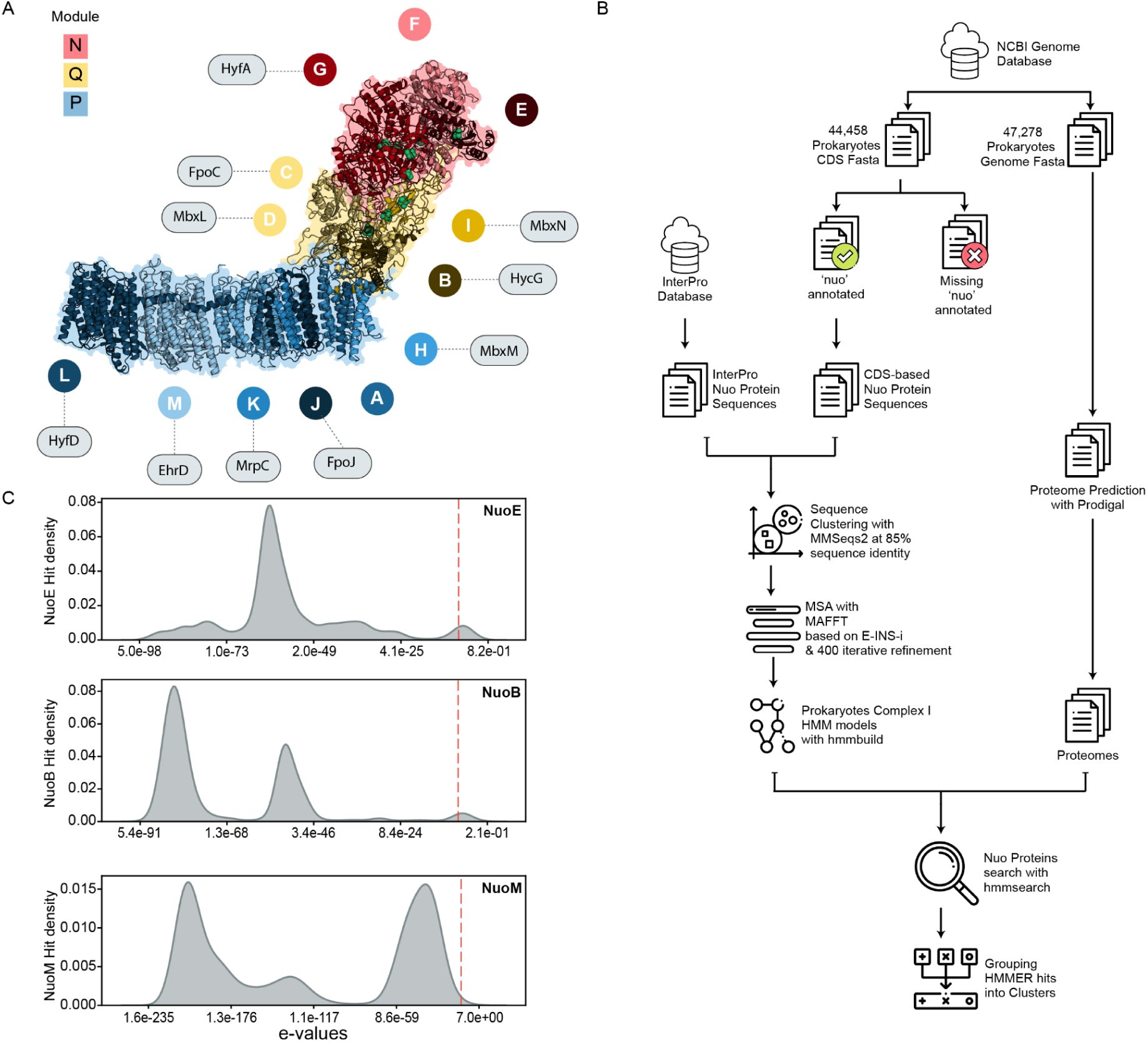
Search for the Nuo subunits in prokaryotic genomes. (**A**) Ribbon diagram of the Complex I structure from *Escherichia coli* (PDB: 7P61) ^18^, color-coded by functional modules: N-module (red) for NADH oxidation, Q-module (yellow) for quinone binding and electron transfer, and P-module (blue) for proton translocation. Labeled circles represent specific Nuo subunits, with homologous proteins from other complexes linked via dotted lines to indicate shared evolutionary origins or functional analogs ^19^. FeS clusters are shown as green spheres. (**B**) Workflow for identifying Nuo subunits in prokaryotic genomes. Sequences from the InterPro database and annotated genomes were clustered using MMseqs2, aligned with MAFFT, and used to build HMM profiles. Nuo-HMMER searches identified Nuo subunit hits, which were grouped into gene clusters ^20–22^ **(C)** KDE-based distributions of Nuo-HMMER hits. The red dashed line indicates one of the generic default thresholds of 1e-7 used for selecting hits in homology-search studies ^13^.

Previous attempts to examine the distribution of Nuo subunits were inadequate due to limited genome sequence data and method precision ^10–13^. The presence of homologs with high sequence and structural similarity confounds the search for valid Nuo subunits. The structural and functional diversity of individual subunits further makes generic search approaches inappropriate. We developed subunit-specific Hidden Markov Model (HMM) profiles for a precise search for Nuo subunits across the entire prokaryotic genome space. We are leveraging a sizeable genome sequence dataset on the NCBI genome database to perform a large-scale distribution analysis. We have: (a) developed a sensitive HMM profile-based sub-unit specific search pipeline for obtaining high confidence hits, (b) obtained taxonomy ranked high-quality genome dataset for 10,608 bacterial and 431 archaeal species for Nuo subunit search, (c) identified Complex I in ten times more bacterial species than previously reported, (d) reported extensive association of prokaryotic lifestyle with Nuo subunits status, (e) identified plasmid-borne Nuo subunits that might have facilitated the distribution of these subunits, and (f) shown prevalence of mitochondrial Complex I accessory subunits in bacterial genomes. We have developed an interactive web application for easy dissemination of knowledge that will enable further experimental efforts in this direction. The alarming rise in resistance against all existing antibiotics has made metabolic pathways an attractive space for the next generation of antimicrobials. A genome-wide understanding of bioenergetic potential is critical in designing optimal therapeutic approaches.

## Results and discussion

### Prokaryotic genome dataset curation

The advent of high-throughput sequencing and associated technological advancements have created a wealth of high-value genomics data. There are more than half a million prokaryotic genome records available on the NCBI Genome database ^14^. We retrieved these genomes and downloaded 47,278 genome sequence fasta files along with their associated 44,458 nucleotide coding sequences (CDS) by applying the filtering criteria of assembly level’complete’ or’chromosome’ (Figure 1B, Supp. Sheet 1). We wanted a clear taxonomic distinction in our study. To obtain the corresponding taxonomy rankings, we used the NCBI taxonomy database to retrieve information for 47,278 genome records through Biopython ^15,16^. Subsequently, we employed TaxonKit v0.16.0 to acquire comprehensive, hierarchical taxonomic information for each taxid included in the genome metadata ^17^. After this initial retrieval and processing, our dataset comprised 10,608 unique bacterial species, represented by their 46,702 strain genomes, and 431 archaeal species, represented by 576 strain genomes.

The selected genomes were further refined based on specific taxonomic criteria to ensure consistency and reliability in downstream analyses. Genomes without designated phylum and genus classifications were excluded from the dataset, as were those marked with “Candidatus” status. This filtering step helped to ensure that the genomes included in our study have a consistent taxonomic nomenclature and a uniform structure for detailed analyses.

### Prokaryotic Nuo subunit search using custom HMM profiles

There is a heavy bias towards some bacterial species in research with only five species accounting for more than forty percent of research papers ^23^. In our genome dataset, we observed a similar skewness with approximately thirty percent of genomes representing only ten bacterial species (Supp. Sheet 2). We took a sequence clustering approach to avoid any bias in our profile-building (Figure 1B). We extracted sequences of all fourteen Nuo subunits from the CDS fasta files with existing annotations for *nuo* genes and translated them into corresponding protein sequences. These sequences were supplemented with reviewed Nuo subunit protein sequences retrieved from the InterPro database ^24^. We clustered these datasets with a threshold of 85% identity to eliminate any taxonomic bias while retaining the sequence diversity.

We used the sequences obtained for each subunit to create a position-specific probabilistic model using HMMER ^25^. We, thereby, obtained prokaryotic subunit-specific representative HMM profiles for searching Nuo subunits. To eliminate inconsistencies in gene-calling that may arise from heterogeneous annotation pipelines, we predicted proteomes of all 47,278 prokaryotic genomes using Prodigal ^26,27^. We, then, performed a search for Nuo subunits by applying our custom HMM profiles to these proteomes. We obtained approximately one million cumulative hits for Nuo subunits of which 6649 hits came from plasmids.

We examined the distribution of HMMER-derived e-values for each Nuo subunit hit, employing kernel density estimates (KDE) to visualize how confidently each hit aligns with the search profiles ^28^. We observed a multimodal distribution of hits for each Nuo subunit (Figure 1C, Supp. Figure 1). Modes with lower e-values can be assumed to consist of high-confidence hits; however, we need further support to set the e-value cutoff as the default threshold could have included several other modes as well ^13^.

### Nuo-HMMER hits clustering to identify high-confidence hits

The extensive sharing of ancestry among functionally similar proteins results in sequence and structure overlap that can pose the risk of false positive calls and Nuo subunit hits are particularly susceptible to such anomalies. Prokaryotic genomes tend to cluster genes belonging to the same physiological functions in neighboring genomic loci on chromosome ^29,30^. We chose to rely on the convergence in the physiological function and consequent clustering of the Nuo subunits to obtain high-confidence hits ^31^. We developed clustering criteria for the Nuo subunit hits based on their respective intergenic distances across all species. Furthermore, for the hits to be clustered together they were required to be present on the same strand. We obtained two types of clusters: (a) Nuo14 cluster containing all fourteen subunits and (b) Nuo-Partial cluster lacking one or more subunit(s). As expected, we observed an initial linear increase in the number of species with Nuo14 cluster hits as we relaxed the intergenic distance criteria for cluster calling, whereas a more stringent cutoff pushed many clusters into the Nuo-Partial category (Figure 2A). However, the increase in the number of species started to flatten beyond 100 base distance and almost plateaued at an intergenic distance of 250 bases. Thus, we put a limit of a maximum of 250 base distance between hits to be grouped in a cluster (Figure 2B). It is noteworthy that our clustering parameter may or may not follow the classical operon definition due to a lack of empirical data.

**Figure 2:**
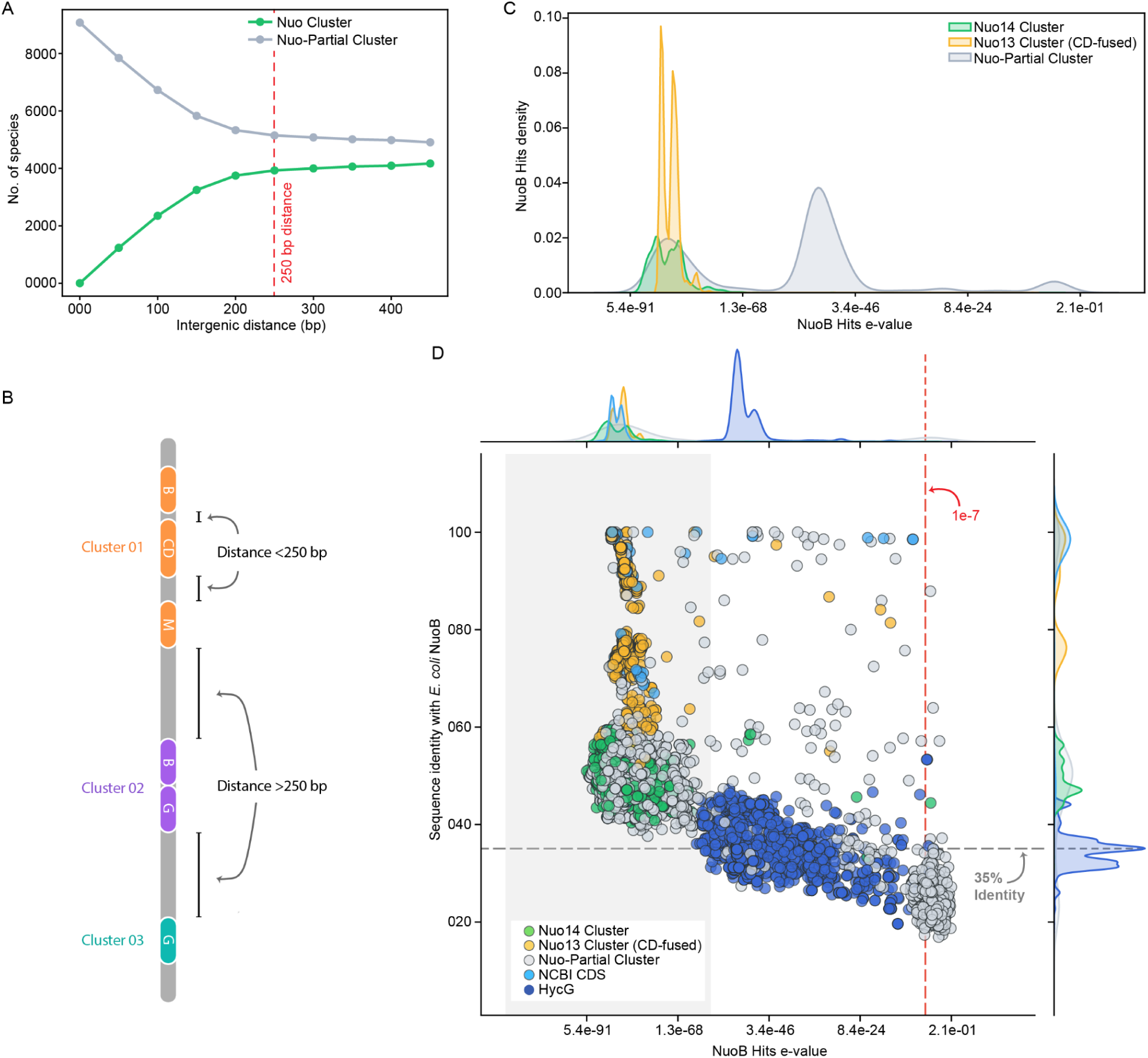
Identification of high confidence Nuo subunit hits and characterization of NuoB subunit hits. **(A)** Influence of maximum allowed intergenic distance for clustering on the number of species assigned to the Nuo14 cluster (green) and Nuo-Partial cluster (gray). The 250 bp threshold (red dashed line) was selected to optimize gene grouping into meaningful clusters. **(B)** Schematic representation of hits clustering based on intergenic distance. Protein subunit hits (their respective genes) within 250 bp are assigned to the same cluster (Cluster 01, orange), while those separated by more than 250 bp are assigned to distinct clusters (e.g., Cluster 02, purple, and Cluster 03, teal). **(C)** KDE-based distributions of NuoB subunit search hits’ e-values. Hits belonging to the Nuo14 cluster and Nuo13 cluster (with fused CD) are highlighted in green and yellow, respectively. **(D)** Scatter plot of NuoB subunit hits showing sequence identity with *E. coli* NuoB sequence (*UniProt*: P0AFC7) against e-values. NuoB subunit hits are color-coded as explained in the inset. The hits with existing NuoB annotation on NCBI are colored in light blue. The hits classified as HycG by COG are highlighted in dark blue. The shaded region (light gray) represents hits filtered based on the e-value threshold obtained from Nuo hits clusters analysis. Marginal density plots illustrate the distribution of e-values (top) and sequence identity (right).

We categorized the hits on the e-value KDE plots according to their clustering pattern. We have segregated the Nuo cluster into (i) the Nuo14 cluster and (ii) the Nuo13 cluster with fused C and D subunits (Figure 2C and Supp. Figure 2). We observed significantly lower e-values for Nuo hits found in clusters and these hits aligned with the left-most mode on the KDE plot. This distinction in segregation allowed us to define e-value cutoffs specific to each Nuo subunit. The presence of fused variants (e.g., NuoCD) underscores the pipeline’s ability to handle non-traditional gene structures. Even in cases where canonical gene boundaries are altered, the established e-value thresholds remain effective, indicating that HMM profiles are robust tools for recognizing modified arrangements. Overall, the lower e-values and an empirical setting of the cut-off allowed us to remove low-confidence hits (Supp. Table 1).

We further analyzed the hits by plotting their protein length against the e-value. It was gratifying to note that the Nuo hits arranged in clusters showed a length distribution within the range coinciding with UniProt for the corresponding subunits (Supp. Figure 3) ^32^. This observation allowed us to eliminate outliers based on their unusual protein length. From a broader perspective, the e-value-based filtering strategy complements the earlier genomic context and clustering analyses. While the operon structure—defined by parameters like intergenic distance and strand orientation—provides a genomic framework, the KDE distributions offer a sequence-level measure of confidence.

We also compared the protein sequence identity of the hits with the corresponding subunit from *Escherichia coli*. Although we did not use sequence identity as a filtering criterion, comparing e-values with sequence identity relative to *E. coli* reinforces the robustness of our approach (Figure 2D). Nuo clusters, which yield low e-values under our KDE-based thresholds (grey region in Figure 2D), also tend to exhibit higher sequence identity. Notably, COG classified the majority of the NuoB HMMER hits with poor e-value as the HycG subunit of the formate hydrogenlyase complex ^33^. HycG shares an evolutionary link with NuoB ^34^. The segregation of distant homologs based on our custom e-value thresholding substantiates the sensitivity of our HMMER-driven approach to searching high-confidence Nuo subunits.

### Distribution and coverage of Nuo subunits

Despite the widespread conservation of Complex I, our understanding is restricted to only a few model species studied in the field ^35–38^. An earlier phylogenomics-based study of 970 bacterial genomes reported the presence of complete Complex I in approximately 500 genomes^12^. Our species-level taxonomy ranking obtained for the 46,702 strain genomes allowed us to examine the distribution of Nuo subunits among 10,608 unique bacterial species. The knowledge gap filled by us brought uniformity in the distribution of various Nuo subunits (Figure 3A). The existing NCBI genome annotation for the Nuo subunits shows that complete Complex I is present in approximately 500 species. Remarkably, our search expanded the coverage of complete Complex I by almost ten-fold (Figure 3B). In 1374 species with complete Complex I, subunit D is fused either with subunit(s) C or with both B and C. We found two complete sets of Nuo subunits in 45 species. *Rhodopseudomonas palustris* is reported to harbor two of these complexes to diversify its metabolism ^39^. Another species where we found two complete Complex I was *Acidiphilium cryptum*. *Acidophilum spp*. survive extreme conditions, and an abundance of laterally acquired genes are believed to be responsible for its adaptive metabolic expansion ^40^. The presence of two NADH-driven proton efflux systems could be responsible for their acidophilic nature.

**Figure 3:**
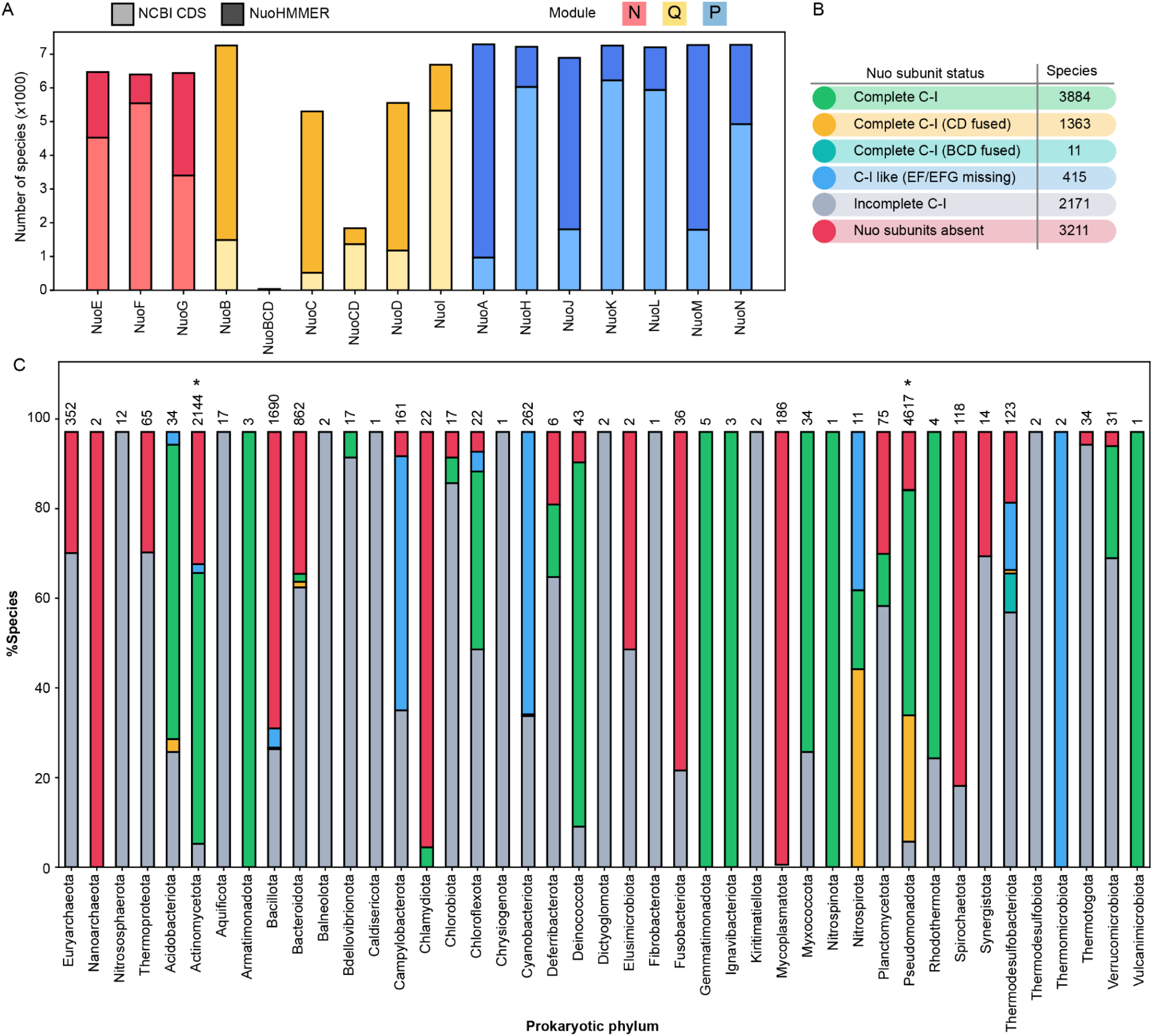
Distribution of Nuo subunits across prokaryotic species. (A) Stacked bar chart showing the number of species with Nuo subunits identified by NuoHMMER and existing NCBI CDS annotations. Bars are color-coded by functional modules: Red (N-module), Yellow (Q-module), and Blue (P-module). (B) Species abundance for Complex I variants based on Nuo subunit status. (C) Percentage distribution of species with different Nuo subunit variations across various phyla. The first four phyla belong to archaea (from left). Bar segments are color-coded to indicate specific subunit variations (refer to panel B). The asterisk represents phyla, where a species shows two different Complex I variants.

A notable group includes 415 species in which this enzyme is classified as “Complex-I like,” because of missing EF or EFG subunits. These subunits constitute the N-module that harbors the FMN cofactor and is responsible for the oxidation of NADH. Such configurations suggest adaptations to specific ecological niches, where the absence of this module may reflect a variation of Complex I to support alternative respiratory strategies. The incomplete Complex I category, comprising 2171 species, represents species where only subsets of Nuo subunits were detected. Some incomplete configurations might have resulted from genomic fragmentation or assembly artifacts. Lastly, 3211 species lacked Nuo subunits entirely, suggesting a reliance on alternative respiratory pathways or metabolic strategies.

Our dataset consisted of genomes belonging to multiple strains of a species. We, therefore, examined the consistency of Complex I (C-I) variants within a species. We observed consistent Complex I variation in all, but one, species (Supp. Sheet 3). *Stenotrophomonas rhizophila* has complete C-I with 14 subunits, whereas one strain has 13 subunits with fused CD. This intra-species variant conservation encouraged us to examine the C-I variant distribution at a higher taxonomic rank. We examined the distribution of C-I variants within every phylum.

We observed the presence of Nuo subunits in three archaeal phyla encompassing seven classes (Figure 3C, Supp. Table 2). Halobacteria and Methanomicrobia showed a C-I-like variant that lacks the N-module. However, our stringent filtering parameter eliminated the subunits I and J from these two classes. Methanomicrobial species *Methanosarcina mazei* complements the N-module deficiency by alternative electron input from the FpoF-mediated oxidation of a lower potential electron carrier F_420_H_2_ ^8,41^. While some of the Nuo hits showed a significant domain similarity with F_420_H_2_ dehydrogenase subunits, the hits obtained for P-module subunits NuoA, NuoM, and NuoN retained their canonical Nuo subunit identity. This subunit arrangement was observed in thirteen species of Methanomicrobia suggesting a widespread use of cofactor F_420_H_2_ as the electron donor in this class. These observations strengthen the proposed course of evolution of Complex I, where P-Q modules are initially believed to have assembled, and later, bacteria acquire N-modules to exploit higher potential electron carriers. Notably, the coupling efficiencies achieved using NADH as a cofactor are significantly higher compared to F420 ^41^.

Bacterial phyla showed a varied distribution pattern for the C-I variant (Figure 3C). Actinomycetota and Pseudomonadota, making up around 64% of the total bacterial species in our dataset, showed several variants of complex-I with a higher prevalence of the complete C-I. In Acidobacteriota, most species showed complete C-I which may be related to their acid tolerance given the hypothesis that pH homeostasis was the selection pressure for the evolution of ATP-independent proton transporters ^4,42^. We found several bacterial phyla where Nuo subunits were mostly lacking. These phyla include Bacillota (with 16% of the total bacterial species in our search), Mycoplasmatota, Spirochaetota, Chlamydiota, Fusobacteriota, etc. Mycoplasmotota and Spirochaetota consist of bacterial species that are obligately dependent on hosts ^43^. *Mycoplasma pneumoniae* is known to lack a functional respiratory chain and rely on organic acid fermentation for ATP supply ^44^. Similarly, species from Chlamydiota are also reported to depend on the host for their metabolic energy requirements ^45,46^.

We have an interesting observation from Thermodesulfobacteriota which are believed to be anaerobic ^47^. This phylum has ten distinct classes, out of which four classes (Desulfuromonadia, Desulfobacteria, Desulfovibrionia, and Desulfarculia) showed a distinct complete C-I variant consisting of fused BCD subunits (Supp. Table 3). Desulfobacterial species inhabit cyanobacterial microbial mats resulting in toxic exposure to oxygen and their oxidative stress defense strategies involve reduction of oxygen using Cytochrome *bd* oxidases ^48^. While this oxidase is suggested to mitigate the load of reactive oxygen species, Cytochrome *bd* oxidases are often part of aerobic ETS ^49^. Notably, Thermosulfobacteria can survive suboxic conditions, and our observation of complete Complex I in this class raises a possibility of adaptive oxic energy metabolism ^50^.

### Association of C-I variants with taxa and metabolic lifestyles

We further focussed on the association of complete C-I variants with various prokaryotic taxa (Figure 4A). We observed five major phyletic clades. In clade 2, we observed a C-I-like variant to form a monophyletic distribution of Campylobacterota suggesting an overall sequence diversity compared to other variants. We observed two distinct sub-clades in clade 1. One of these two subclades consists of C-I-like variants coming from ancient bacterial phyla Bacillota and Cyanobacteriota. Cyanobacteriota is believed to have functional Complex I but the identity of the N-module supplement is under investigation ^8^. The majority of the clades have a mix of split (Nuo subunits not clustered together) and single cluster Complex I, however, the subclades occupied by Actinomycetota in clade 3 and Gammaproteobacteria and Betaproteobacteria in clade 4 are dominated by single cluster Complex I (Supp. Figure 4). The clades 4 and 5 showed polyphyletic distribution with species from Gammaproteobacteria and Alphaproteobacteria split on them. Both these clades consist of complete C-I variants with clade 5 populated variants with fused C and D subunits. We looked at the evolutionary history of the relevant C-I variants after Gammaproteobacterial species split into two separate clades. In order to achieve this, we overlaid C-I variant data onto a rooted phylogenetic tree of Pseudomonadota species (Supp. Figures 5 and 6). The phylogenomic timeline of proteobacterial divergence is reflected in the tree structure. Alphaproteobacteria first appeared around 1.9 billion years ago, followed by Betaproteobacteria and Gammaproteobacteria as the most derived subgroups ^51,52^. Upon layering the C-I variant information on this species tree, we observed a distinctive pattern. The evolutionary older species showed the presence of 14 subunits complete C-I, whereas modern species have 13 subunits C-I variant with C and D subunits fused. Functional synergy is often a cause of gene fusion, and modern variants with fused subunits support the evolutionary convergence of distinct proteins to achieve a collective physiological function ^53^.

**Figure 4:**
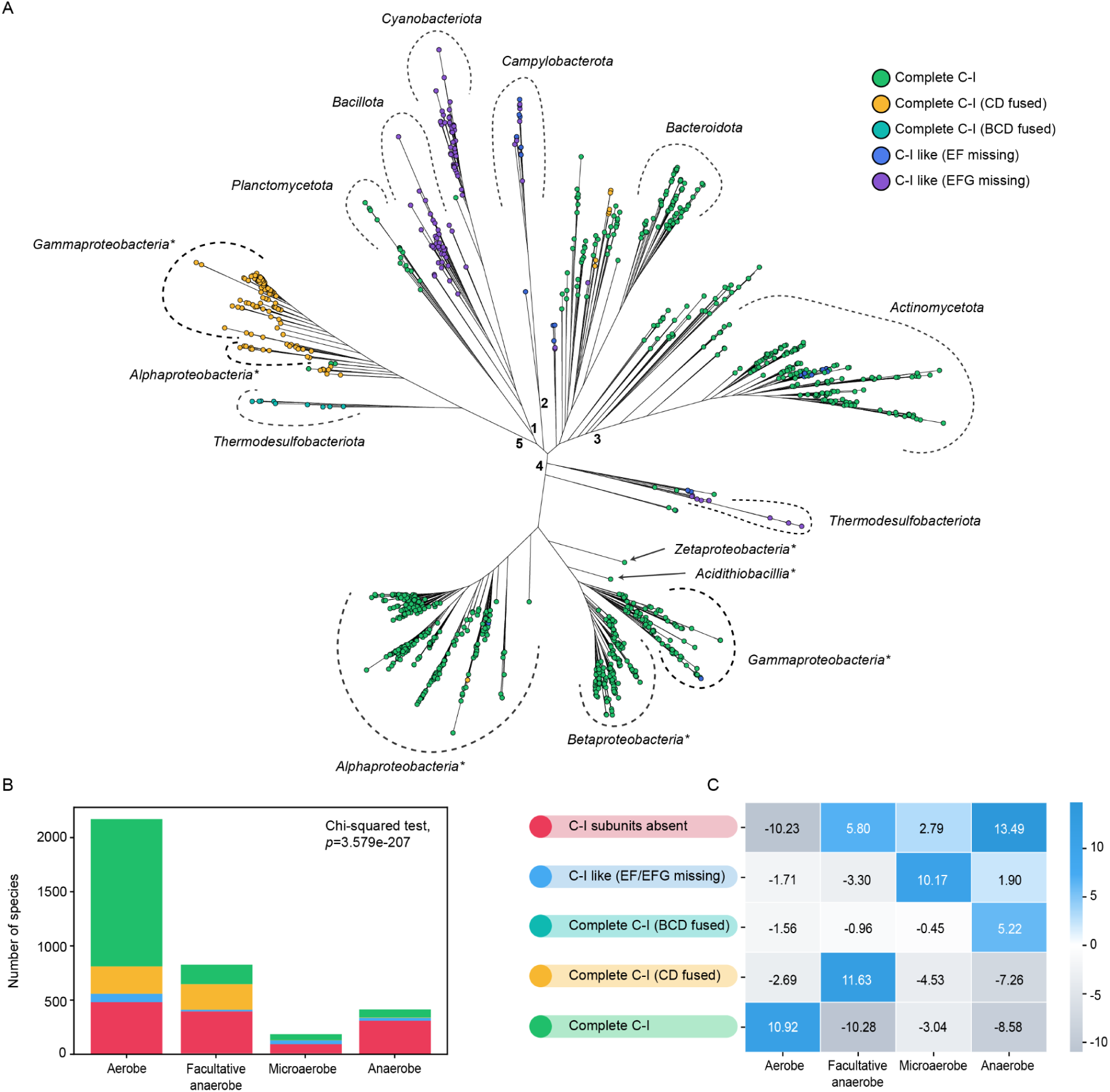
Association of C-I variants with taxa and metabolic lifestyles. (A) The phylogenetic tree is based on concatenated sequences of Nuo subunits from one species per genus. The C-I variations have been shown by color-filled circles. Bacterial taxonomic phyla and classes (marked with an asterisk) have been highlighted using curved dashed lines. (B) Distribution of species with various metabolic lifestyles (Aerobe, Anaerobe, Facultative Anaerobe, and Microaerobe) based on C-I variations. The statistical significance of the relationship between oxygen tolerance and C-I status was assessed using a chi-square test. (C) Heatmap showing residuals from the chi-square test, representing deviations from expected distributions of C-I variants for a given metabolic lifestyle. Positive values (blue shades) indicate an association, while negative values (gray shades) indicate dissociation.

**Figure 5:**
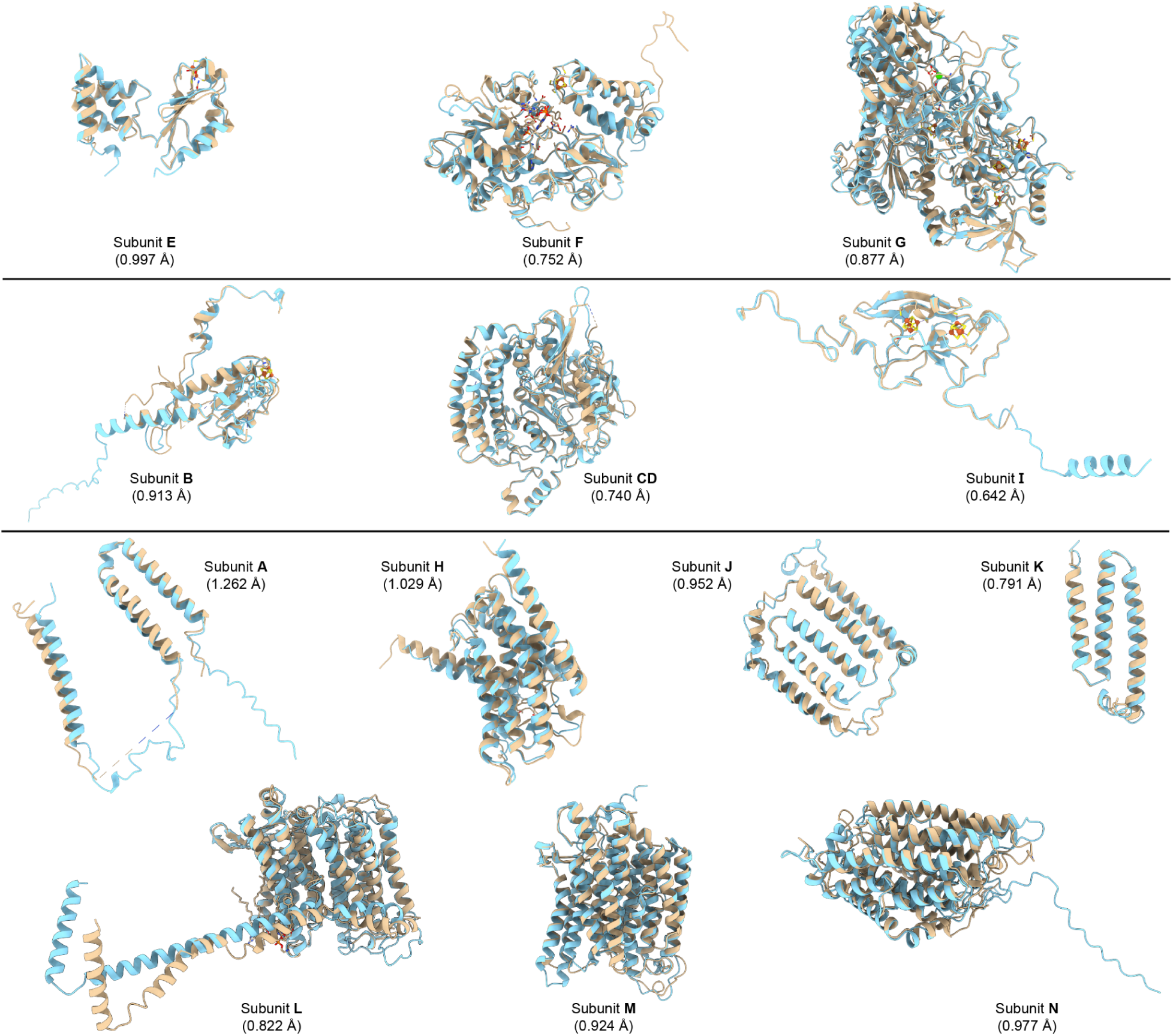
Structural comparison of Nuo subunits from the plasmid. The alignment of cryoEM structures of *E. coli* Nuo subunits (chromosomal origin, PDB: 7P61, beige) with corresponding AlphaFold3-predicted structure for plasmid-derived hits in *K. saccharivorans* (blue). Their respective RMSD values are shown in brackets.

**Figure 6:**
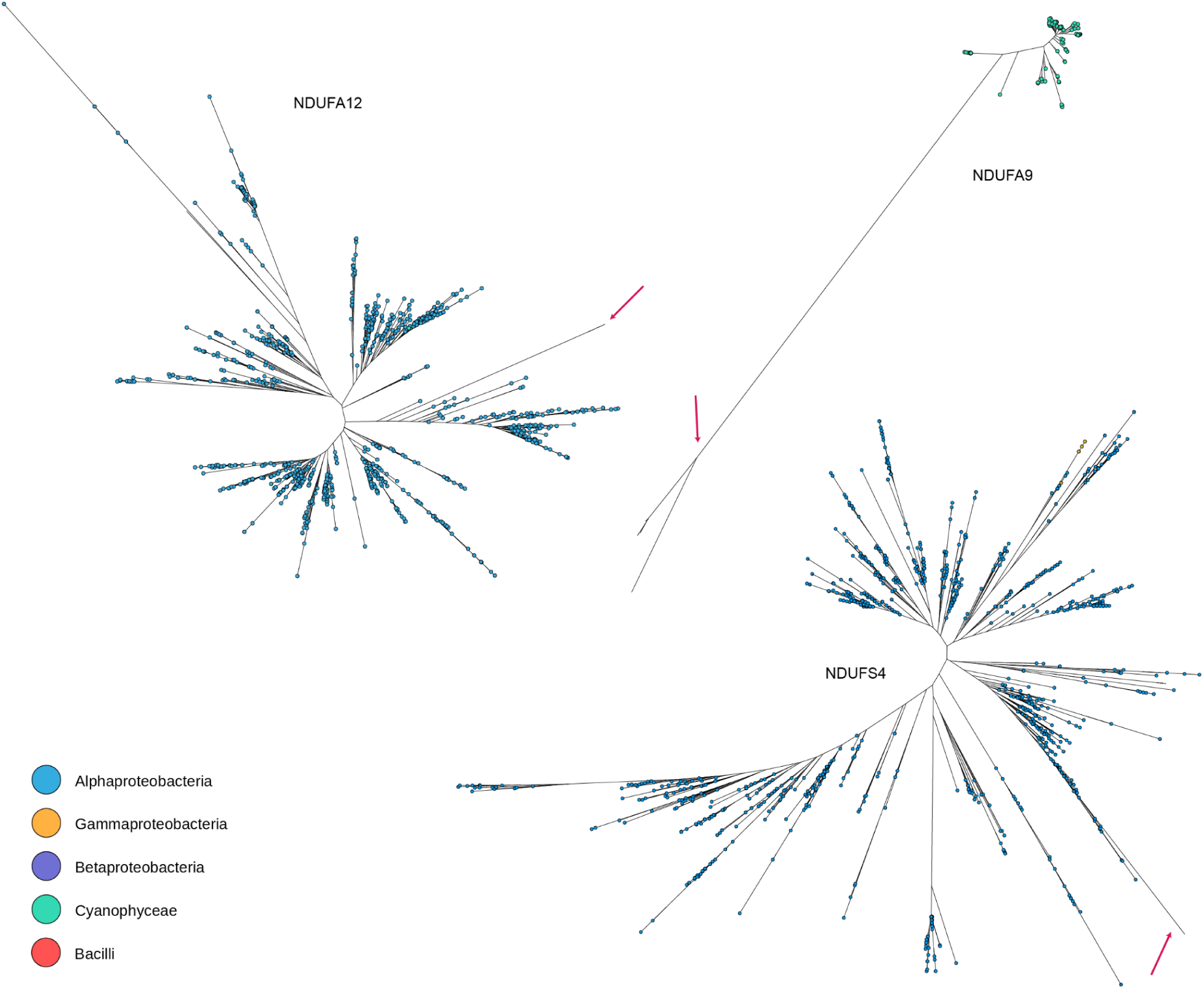
Phylogenetic tree for the Complex I accessory subunits. Maximum Likelihood-based phylogenetic trees were constructed for three accessory subunits (NDUFA12, NDUFA9, and NDUFS4) of Complex I using sequences identified through NuoHMMER workflow. The trees display bacterial species that contain matches for these subunits. Each circle represents a bacterial species, color-coded according to its respective taxonomic class. Eukaryotic sequences are marked with red arrows. Branch lengths indicate evolutionary distance in terms of substitutions per site.

Complex I reduces the respiratory quinones which can then interact with either oxic or anoxic components of the ETS based on the metabolic demand. We were motivated to examine the oxygen-dependent metabolic lifestyle preferences of bacterial species with complete Complex I. We collected lifestyle information from the BacDive database (Supp. Sheet 4) ^54^. We also performed an extensive literature survey to fetch information for species that were missing from the above databases. We observed a significant positive association between the aerobic nature of the bacteria and the presence of complete Complex I (Figure 4B, 4C). Interestingly, obligate aerobes and facultative anaerobes showed distinct C-I variants. A similar association was reported earlier ^12^. It is gratifying that the lifestyle association derived from a smaller dataset of approximately 700 species remains valid for a five times larger dataset. We also observed microaerophilic bacteria to associate strongly with Nuo-like complex I. This group includes *Campylobacter jejuni* which is known to lack a functional N-module and rely on flavodoxin as the source of electrons ^55^. Interestingly, the complete C-I variant with fused BCD subunits showed an association with an anaerobic lifestyle. As discussed in the previous section, the Thermodesulfobacteriota species possessing this complex I variation likely have minimal oxic metabolism.

### Nuo subunits on plasmids

The widespread distribution of Nuo subunits and various C-I variants supports the hypothesis that the subunits evolved independently and came together as units to achieve synergistic outcomes. It is appealing to assume that Nuo subunit genes traveled among prokaryotic species, resulting in the assembly of complete Complex I ^56^. Typically, such gene transfer events are mediated by plasmids, which carry genes that assist their host in acquiring adaptive traits ^30^.

Our dataset allowed us to distinguish the Nuo subunit hits coming from chromosomes and plasmids. We noted that Nuo subunits were located on plasmids in 57 species (Supp. Sheet 5). While in most cases, only a partial set of Nuo subunits was obtained, 17 species showed a complete set of Nuo subunits on plasmids (Supp. Table 4). Interestingly, Nuo subunits were exclusively located on plasmids in five bacterial species. This genomic distribution of Nuo subunits has never been reported; therefore, we performed further validation for this observation. We picked the plasmid-borne Nuo subunit sequences from *Komagataeibacter saccharivorans* and built protein structures for individual subunits using AlphaFold3 ^57^. Then we examined their structural conservation with cryo-EM derived structures of chromosomally located *E. coli* Nuo subunits. We obtained very high structural conservation between corresponding subunits. The presence of Nuo subunits, and in a few instances complete C-I, on plasmids indicates a potential mechanism of intra-and inter-species transfer of corresponding genes.

### Mitochondrial Complex I accessory subunits in prokaryotes

The evolutionary assembly of multiple subunits to form the complete complex I has continued in eukaryotes and mitochondrial complex I has acquired additional subunits to form a 45-subunit complex. These additional subunits are accessory to the core subunits, but their presence is critical for the assembly and function of human mitochondrial complex I ^58^.

Interestingly, while bacterial Complex I with fourteen subunits is fully functional, recent studies have shown a few accessory subunits in some bacteria. The cryo-EM structure of *Mycobacterium smegmatis* complex I showed subunit A9 as part of the complex whereas three accessory subunits (A12, S4, and S6) were reported from *Paracoccus denitrificans* ^37,38^.

We, therefore, followed our workflow to probe the distribution of such accessory subunits in prokaryotes. In this case, the HMM profiles for all 31 mitochondrial accessory subunits were built using corresponding sequences from the InterPro database which were, as expected, dominated by eukaryotic sequences. We observed hits for A9, A12, and S4 accessory subunits in 1402 bacterial species out of which 1279 species belong to Alphaproteobacteria (Supp. Sheet 6, Supp. Figure 7). Notably, we have found the complete C-I variant in these alphaproteobacterial species. This close association of accessory subunits with complete Complex I suggests their functional involvement. Notably, both Rhodobacterales and Rickettsiales, which are argued to be closer to the mitochondrial complex I evolution, belong to this class ^59–61^. The subunit A9 is absent across all Alphaproteobacterial species and was found localized in 115 species of Cyanophyceae. These observations suggest a more distributed evolution of Complex I accessory subunits in prokaryotes, which would have converged later during the endosymbiont event.

The assembly of various proteins with distinct biochemical functions for the evolution of Complex I is a marvel of molecular innovation. Such complexation is driven by selection pressure on assembly intermediates, and evolution selects for protein complexes that assemble via ordered pathways ^62^. An appealing immediate prospect of our study would be to map the Nuo subunits identified in this work onto the bacterial tree of life to trace the evolutionary course of subunits’ assembly. Active efforts are underway to exploit bacterial energy metabolism as the target space for antibiotic development ^63,64^. Given its absence from mitochondrial energetics, the non-proton-pumping alternate to complex-I, type II NADH: quinone oxidoreductase is considered an effective target ^65^. However, the success of such approaches critically depends on detailed and precise knowledge of the pathogen’s metabolic capacity. We have developed a web application (prokcomplexone.streamlit.app) on the Streamlit platform to provide an interactive platform for researchers to explore the results of our comprehensive study on the distribution of Nuo subunits and diversity of Complex I across prokaryotic genomes. This WebApp will bridge the gap between experimental and computational analyses and potentiate translational efforts.

## Materials and Methods

### Data Acquisition

As of September 5, 2024, the NCBI Genome database contained 729,729 prokaryotic genome records. For this study, we selected 47,278 genomes classified as either “Complete Genomes” or “Chromosomes.” This dataset includes 46,702 bacterial genomes and 576 archaeal genomes (Supp. Sheet 1). Taxonomic rankings were retrieved using TaxonKit v0.14.0 ^17^ via NCBI TaxIDs, ensuring consistent species-level resolution for downstream analyses. Our genome dataset includes 11,039 prokaryotic species.

### Nuo subunit-specific HMMER profiles construction

Annotated *nuo* subunit sequences were extracted from CDS FASTA files, translated into corresponding protein sequences, and supplemented with reviewed Nuo subunit sequences from the InterPro database to ensure a comprehensive dataset ^24^. To reduce sequence redundancy, MMseqs2 v13.45111 was used to cluster Nuo proteins at 85% sequence identity ^22^. Multiple sequence alignments were performed using MAFFT v7.490 (E-INS-i strategy), optimized for sequences with multiple conserved domains, and refined over 400 iterative cycles ^21^. Using HMMER v3.4, we built 16 individual HMM profiles (14 for individual subunits and 2 for fused subunits), with hmmbuild, incorporating Laplace priors to enhance statistical robustness ^20^. For each subunit, sequence weighting was disabled (--wnone) to ensure equal representation of all sequences within that subunit’s dataset, while core model definition and fragment handling were adjusted (--symfrac 0.6,--fragthresh 0.3) to balance conservation with sequence variability.

### Systematic identification of Nuo subunits using HMM profiles and filtering

Before HMMER-based subunit detection, proteome prediction for all genome files was performed using Prodigal v2.6.3 ^27^. Subsequently, the predicted proteomes were searched against the NuoHMMER profiles through hmmsearch (HMMER v3.4). Following the HMM search, we analyzed the distribution of e-values for each Nuo subunit hit and employed kernel density estimates (KDE) to assess distribution. To minimize false hits, we implemented a filtering strategy based on three key criteria: hits clustering, KDE-based e-value cutoffs, and protein length validation.

### Metadata curation and associated C-I variant analysis

Metadata on oxygen requirements was retrieved from the BacDive and IMG databases ^27,66^. For species lacking entries in these databases, manual curation from literature sources was performed (Supp. Sheet 4). Oxygen tolerance was categorized into four groups: aerobe, anaerobe, Facultative anaerobe, and microaerobe. To assess the relationship between C-I variants and oxygen tolerance, a chi-squared test of independence was conducted (p < 0.0005). A contingency table was constructed to compare observed frequencies and statistical significance was evaluated.

### C-I variants phylogenetic tree

Phylogenetic analyses were conducted using concatenated protein sequences from 1,196 bacterial species, with one species selected per genus. The selection criteria ensured that each genus included in the dataset possessed Complex I and its variations. Multiple sequence alignment was performed using MAFFT v7.490 ^21^ with the L-INS-i algorithm, optimized for accuracy in large datasets. The alignment was subsequently trimmed using BMGE v1.12 ^67^ to remove poorly aligned and compositionally biased regions. Maximum likelihood (ML) phylogenies were inferred using IQ-TREE v1.6.12 ^68^, with the best-fit substitution model determined by ModelFinder. Branch support was assessed using ultrafast bootstrap approximation (UFBoot, 1000 replicates) and the SH-aLRT test (1000 replicates). The final dataset consisted of 2230 alignment positions, including 2099 parsimony-informative sites, with automatic filtering of gapped and compositionally biased sequences. As an alternative approach, approximately-ML trees were constructed using FastTree v2.1 ^69^ with the WAG + Gamma model, which is optimized for large datasets. The resulting phylogenies were used to investigate the evolutionary relationships of C-I variants across prokaryotic lineages.

### Plasmid-borne Nuo subunit structure prediction

The three-dimensional structures of individual subunits were predicted using AlphaFold3 ^57^. The amino acid sequences of the subunits were used as input for the AlphaFold server. The highest-ranked model based on the predicted Local Distance Difference Test (pLDDT) score was selected for further analysis.

## Mitochondrial Complex I accessory subunits HMM profiles

To identify mitochondrial Complex I accessory subunits in prokaryotic proteomes, we constructed HMM profiles following the same protocol used for Nuo subunits. Instead of extracting sequences from CDS FASTA files, we obtained protein sequences from the NCBI Protein Database and InterPro Database to ensure a comprehensive dataset. To reduce redundancy, sequences were clustered at 85% identity using MMseqs2. Multiple sequence alignments were performed with MAFFT v7.490 (E-INS-i strategy) and refined over 400 iterative cycles. HMM profiles were generated using HMMER v3.4, incorporating Laplace priors for statistical robustness. These HMM profiles were subsequently used for systematic detection of mitochondrial Complex I accessory subunits in prokaryotic proteomes.

### Phylogenetic tree for mitochondrial Complex I accessory subunits identified in prokaryotes

To investigate the phylogenetic relationships among the sequences, multiple sequence alignment (MSA) was conducted using MAFFT with the--localpair option. The resulting alignments were subsequently refined using TrimAl ^70^ with the-automated1 parameter to eliminate poorly aligned or ambiguous regions, ensuring the retention of high-confidence sites for phylogenetic inference. A maximum likelihood (ML) phylogenetic tree was constructed using IQ-TREE, employing the LG+F+I+G4 substitution model. Branch support was evaluated through 1,000 ultrafast bootstrap replicates (-bb 1000) and approximate likelihood-ratio tests (-alrt 1000).

## Supporting information

Supp. Sheet 1: List of genomes retrieved from NCBI genome database

Supp. Sheet 2: List of bacterial species and their genome representation in the analysis

Supp. Sheet 3: List of bacterial species and their associated Complex I variants

Supp. Sheet 4: List of lifestyle information for all species

Supp. Sheet 5: List of bacterial species containing nuo subunit on plasmid

Supp. Sheet 6: List of bacterial species containing mitochondrial Complex I accessory subunits

## Acknowledgments

This work was supported by the DAE-Tata Institute of Fundamental Research Grant (19P0120) and DBT-Ramalingaswami Fellowship (21X432) to Amitesh Anand. We thank Stuti Srivastav, Neha Banwani, Arpita Biswas, Anjali V. Patil, and Amartya Chowdhury from Anand Research Group for their critical review of the manuscript.

## Author contributions

Conceptualization and Supervision: A.A.; Methods: A.S. and Ab.A.; Analysis: A.S., Ab.A. D.M.P., S.K., and A.A.; Manuscript writing A.A., A.S., and S.K.

## Data and code availability

The data used in this paper are publicly available. The processed data and scripts used in the manuscript have been uploaded to and are available at the GitHub repository: https://github.com/Anand-Research-Group/complex-I.

## Notes

### Competing Interest Statement

The authors have declared no competing interest.

