## Supplementary figures and tables for "Hidden Markov Model-Based Prokaryotic Genome Space Mining Reveals the Widespread Pervasiveness of Complex I and Its Potential Evolutionary Scheme"

### Content

**Figures:**

**Supp. Figure 1:** KDE-based distributions of Nuo-HMMER hits.

**Supp. Figure 2:** KDE-based distributions of e-values of Nuo subunit search hits.

**Supp. Figure 3:** Nuo hits distribution by plotting their protein length against the e-value.

**Supp. Figure 4:** Phylogenetic tree of Nuo subunits, representing one species per genus.

**Supp. Figure 5:** Phylogenetic tree of Pseudomonadota

**Supp. Figure 6:** Phylogenetic tree of Pseudomonadota annotated with Complex-I variants.

**Supp. Figure 7:** Phylogenetic tree of Pseudomonadota annotated with accessory subunits of Complex-I.

**Tables:**

**Supp. Table 1:** Custom e-value cutoff for individual Nuo subunits.

**Supp. Table 2:** Distribution of Complex I variants in archaeal phyla

**Supp. Table 3:** Distribution of Complex I variants in Thermodesulfobacteriota

**Supp. Table 4:** List of species with a complete set of Nuo subunits on plasmids

**List of supplementary sheets:**

**Supp. Sheet 1:** List of genomes retrieved from NCBI genome database

**Supp. Sheet 2:** List of bacterial species and their genome representation in the analysis

**Supp. Sheet 3:** List of bacterial species and their associated Complex I variants

**Supp. Sheet 4:** List of lifestyle information for all species

**Supp. Sheet 5:** List of bacterial species containing nuo subunit on plasmid

**Supp. Sheet 6:** List of bacterial species containing mitochondrial Complex I accessory subunits

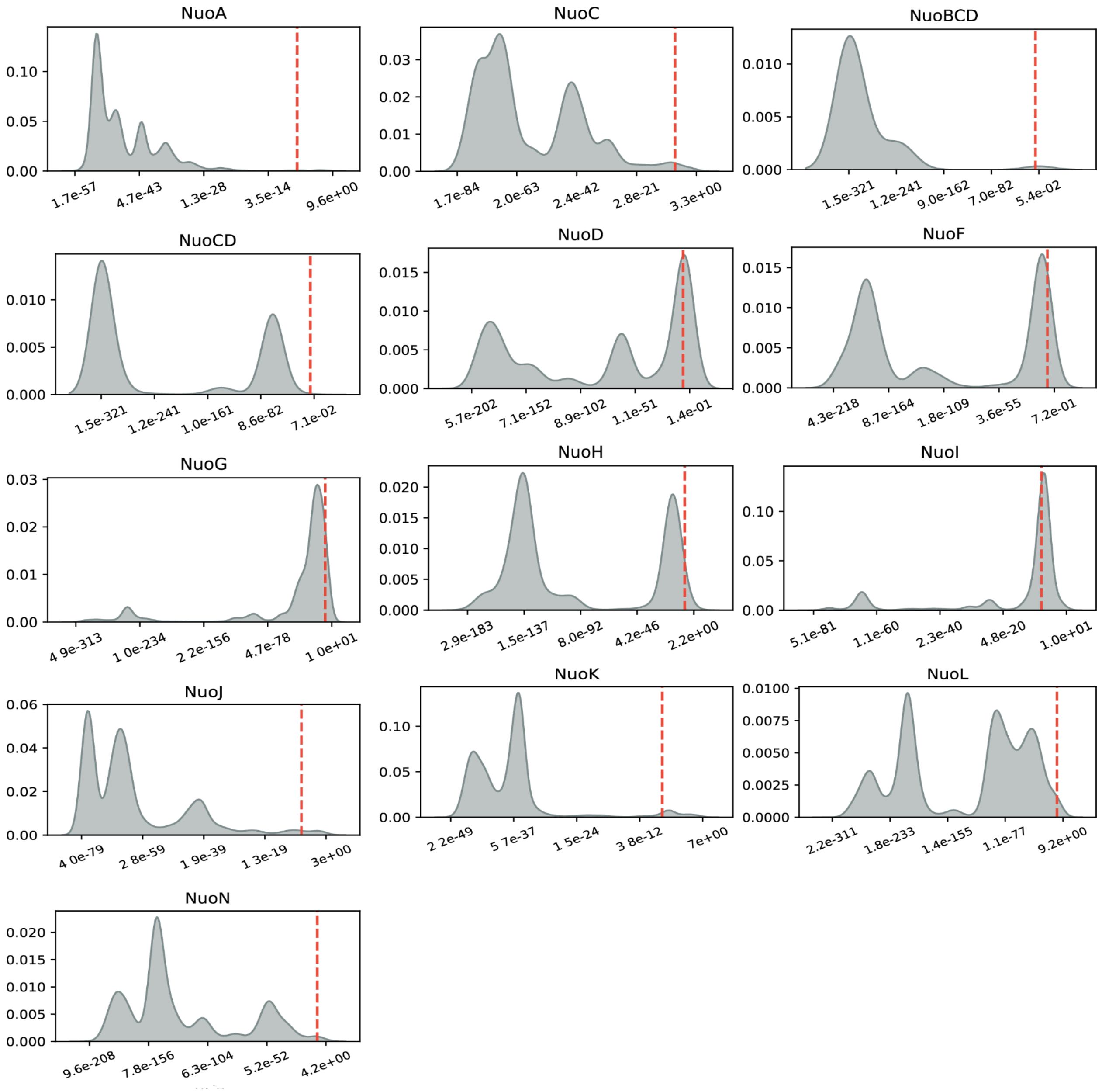


**Supp. Figure 1:**Kernel Density Estimation (KDE)-based distributions of Nuo-HMMER hits for each Nuo subunit. The x-axis represents e-values, while the y-axis denotes the density of hits. The red dashed line marks the commonly used default threshold of 1e-7 for hit selection in homology-based searches.

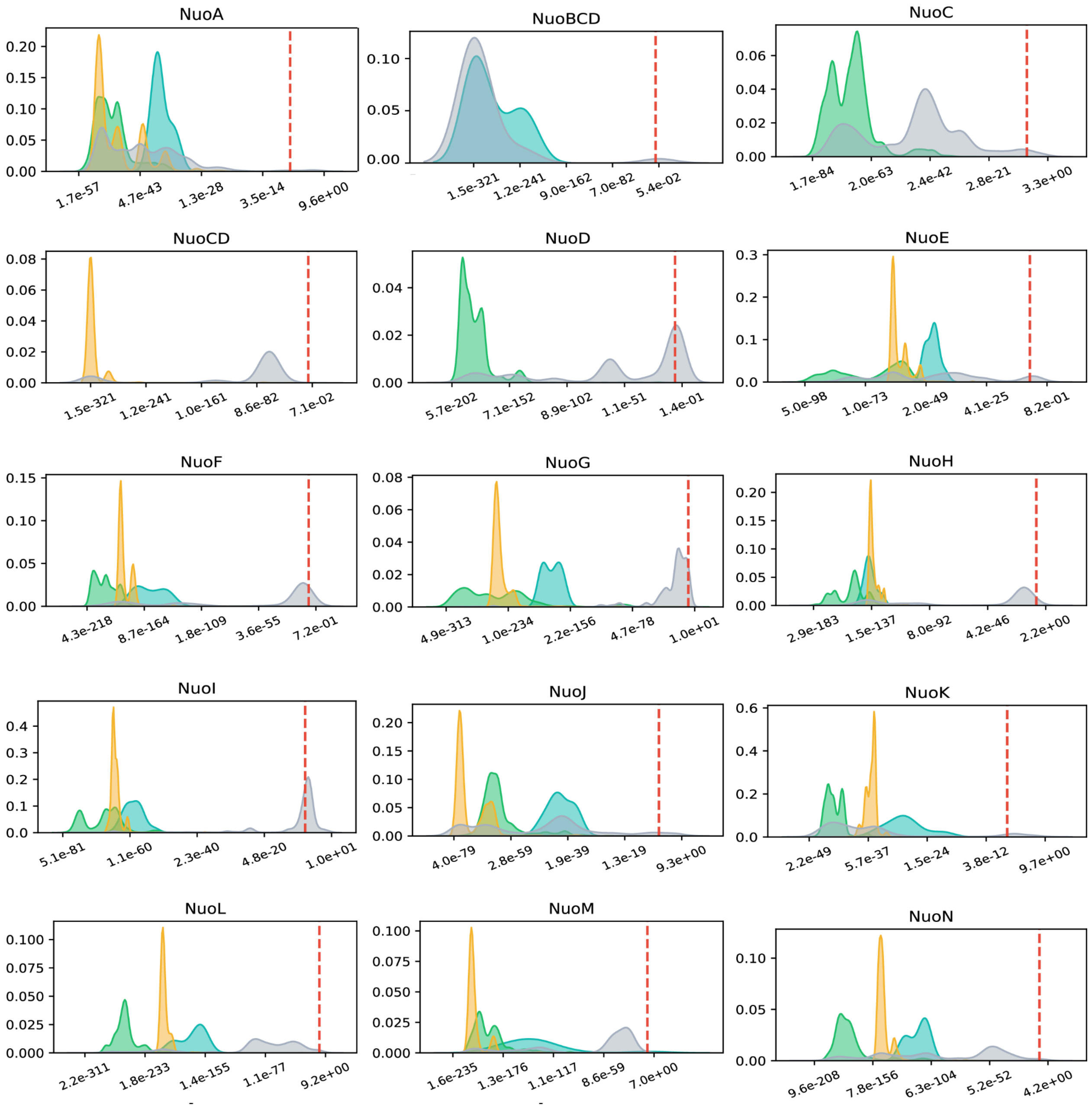


**Supp. Figure 2:** Kernel Density Estimation (KDE)-based distributions of Nuo subunits search hits' e-values. Hits belonging to the Nuo14 cluster and Nuo13 cluster (with fused CD) are highlighted in green and yellow, respectively. The x-axis represents e-values, while the y-axis denotes the density of hits. The red dashed line marks the commonly used default threshold of 1e-7 for hit selection in homology-based searches.

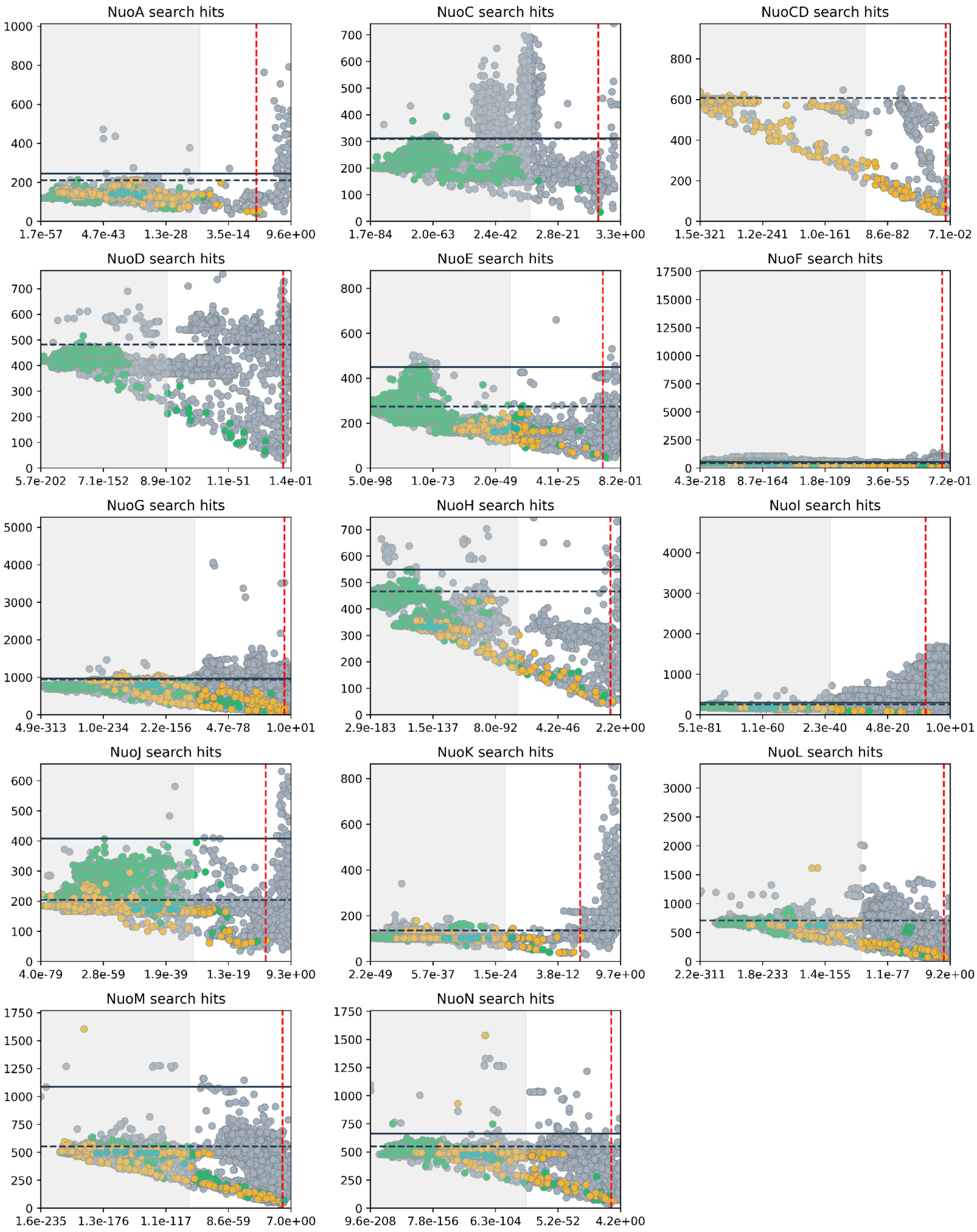


**Supp. Figure 3:** Nuo hits distribution by plotting their protein length against the e-value.Hits belonging to the Nuo14 cluster and Nuo13 cluster (with fused CD) are highlighted in green and yellow, respectively. The x-axis represents e-values, while the y-axis denotes the protein length of hits. The black dashed line is the Uniprot maximum length for a given subunit. The black solid line is the threshold chosen to remove false hits. The red dashed line marks the commonly used default threshold of 1e-7 for hit selection in homology-based searches. The gray region is a subunit-specific e-value cutoff region.

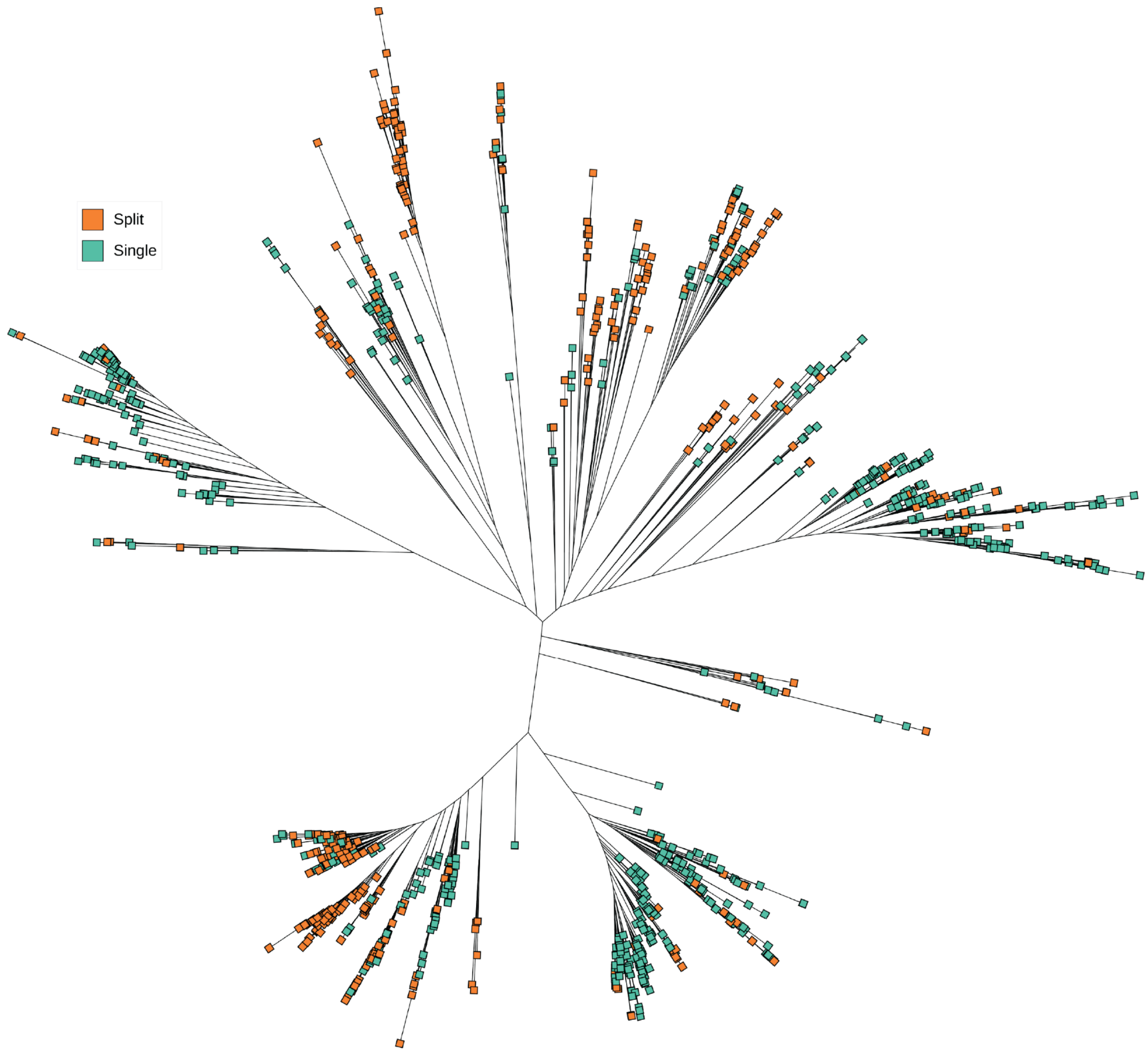


**Supplementary Figure 4:** Phylogenetic tree constructed from concatenated sequences of Nuo subunits, representing one species per genus. The arrangement of Nuo subunits is indicated by color-filled circles. ‘Single’ denotes all *nuo* genes clustered together within a distance of <250 bases, while ‘Split’ indicates the presence of multiple clusters where *nuo* genes are separated by >250 bases.

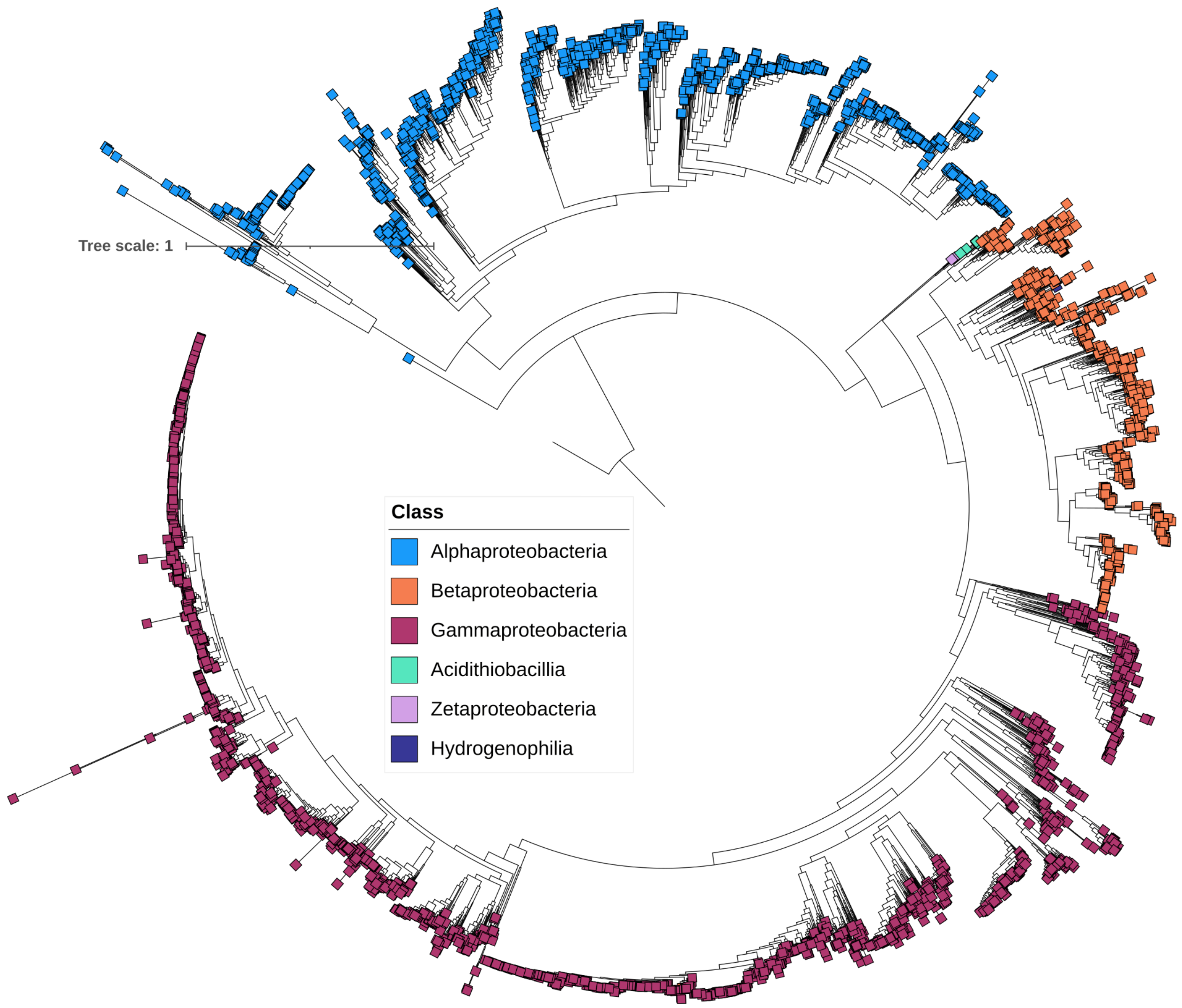


**Supplementary Figure 5:** Pseudomonadota (Proteobacteria) circular phylogenetic tree built with GToTree, rooted with *Acidobacterium capsulatum* from the phylum Acidobacteriota. Branch lengths show the relative divergence of species by representing evolutionary distances. According to the legend, Pseudomonadota's taxonomic classes are colour-coded. A relative indicator of evolutionary divergence is shown by the scale bar.

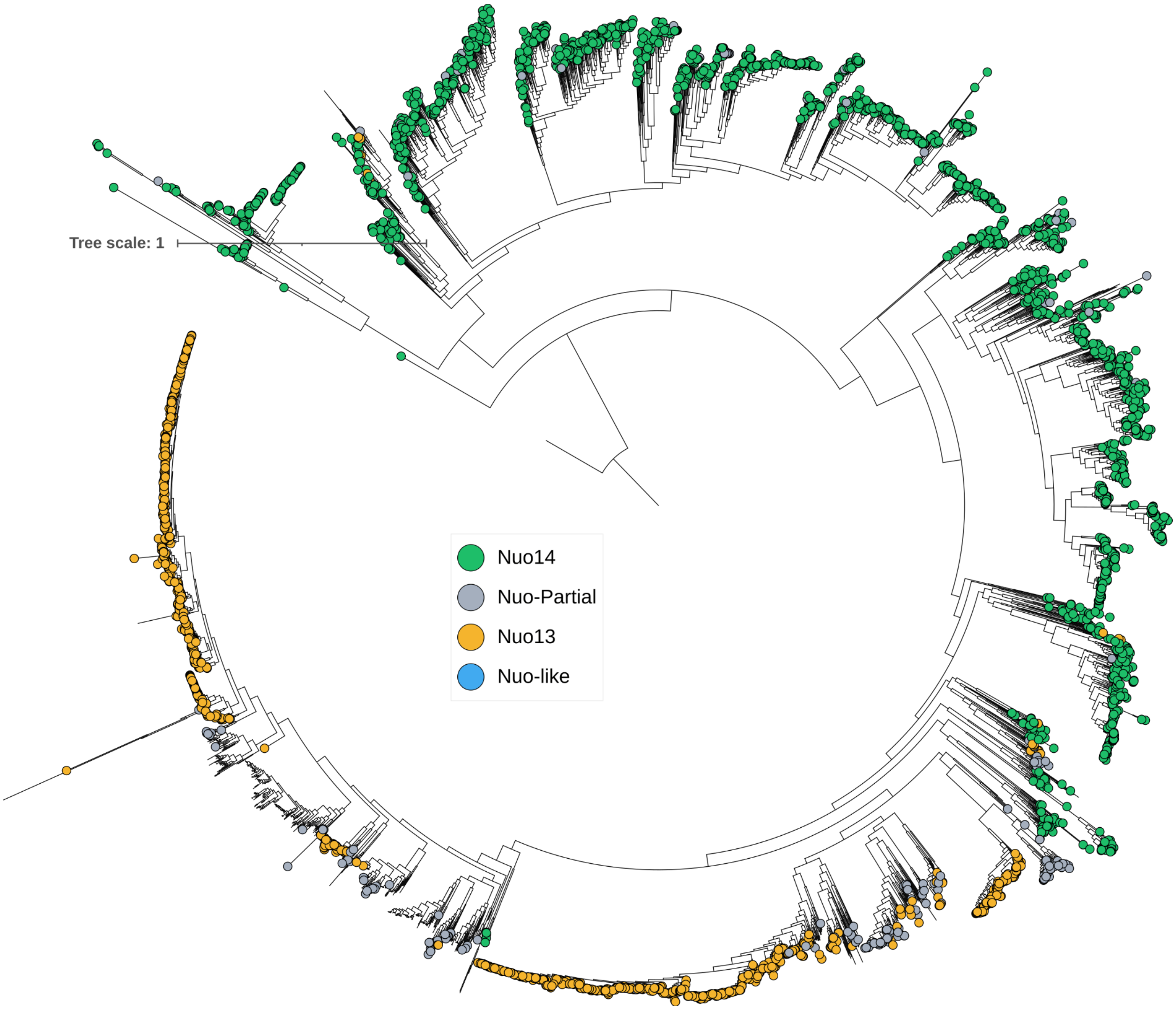


**Supplementary Figure 6:** Pseudomonadota (Proteinobacteria) circular phylogenetic tree annotated with Complex I variations. Branch lengths show species difference by corresponding with evolutionary distances. As the legend notes, complex I versions are color-coded. The scale bar provides a relative indication of evolutionary separation.

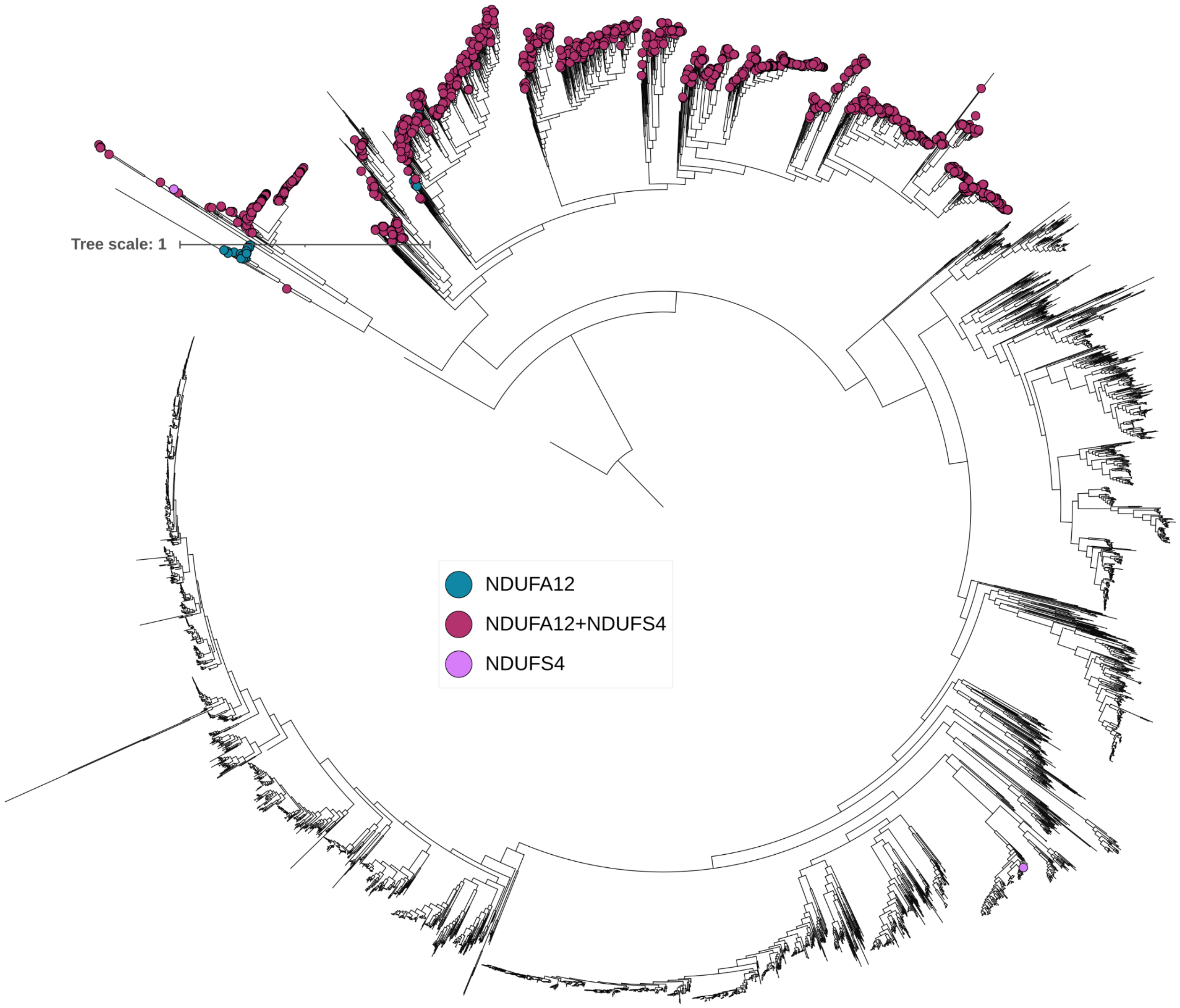


**Supplementary Figure 7:** Pseudomonadota's (Proteobacteria) circular phylogenetic tree with Complex I accessory subunits annotated. Branch lengths show species separation by corresponding to evolutionary distances. The legend indicates the color-coded presence of accessory subunits. A relative indicator of evolutionary divergence is shown by the scale bar.

**Supp. Table 1:** Nuo subunit specific e-value Cutoff

| Nuo subunit | log10(e-value) | e-value |
|---|---|---|
| NuoA | -20 | 1.00E-20 |
| NuoB | -60 | 1.00E-60 |
| NuoBCD | -200 | 1.00E-200 |
| NuoC | -30 | 1.00E-30 |
| NuoCD | -110 | 1.00E-110 |
| NuoD | -100 | 1.00E-100 |
| NuoE | -43 | 1.00E-43 |
| NuoF | -74 | 1.00E-74 |
| NuoG | -120 | 1.00E-120 |
| NuoH | -75 | 1.00E-75 |
| NuoI | -38 | 1.00E-38 |
| NuoJ | -30 | 1.00E-30 |
| NuoK | -22 | 1.00E-22 |
| NuoL | -110 | 1.00E-110 |
| NuoM | -95 | 1.00E-95 |
| NuoN | -78 | 1.00E-78 |

**Supp. Table 2:** Distribution of Complex I variants in Archaea

| Phylum | Class | Variant |
|---|---|---|
| Euryarchaeota | Methanomicrobia, Halobacteria, Methanobacteria, Thermococci, Archaeoglobi | Incomplete C-I |
| Nitrososphaerota | Nitrososphaeria | Incomplete C-I |
| Thermoproteota | Thermoprotei | Incomplete C-I |

**Supp. Table 3:** Class wise distribution of Complex I variants in Thermodesulfobacteriota phylum

| Class | Complete C-I (BCD Fused) | Complete C-I (CD Fused) | C-I like (EF/EFG missing) |
|---|---|---|---|
| Desulfarculia | *Desulfoferula mesophila* | | |
| Desulfobaccia | | | |
| Desulfobacteria | *Desulfonema limicola*; *Desulfococcus multivorans*; *Desulfosarcina alkanivorans*; | | |
| Desulfobulbia | | uncultured *Desulfobulbus* sp. | *Desulfobulbus oralis* |
| Desulfomonilia | | | |
| Desulfovibrionia | | | *Desulfovibrio desulfuricans*; *Desulfovibrio* sp. G11 |
| Desulfuromonadia | *Syntrophotalea carbinolica* *Desulfuromonas soudanensis* *Syntrophotalea acetylenica* *Syntrophotalea acetylenivorans* *Desulfuromonas versatilis* *Geoalkalibacter halelectricus* *Desulfuromonas sp. AOP6* | | *Desulfuromonas acetoxidans* |
| Syntrophia | | | |
| Syntrophobacteria | | | |
| Thermodesulfobacteria | | | *Thermodesulfobacterium commune*; *Thermodesulfobacterium* sp. TA1; *Caldimicrobium thiodismutans*; *Thermosulfuriphilus ammonigenes*; *Thermodesulfatator indicus;* *Thermodesulfobacterium geofontis*; |

| | | | |
|---|---|---|---|
| | | | *Thermosulfurimonas marina* |

**Supp. Table 4**: List of species showed a complete set of Nuo subunits on plasmids.

| **Species** | **Accession** | **Variation** |
|---|---|---|
| *Salmonella enterica* | NZ_CP148876.1 | Complete C-I (CD Fused) |
| *Escherichia coli* | NZ_CP141089.1 | Complete C-I (CD Fused) |
| *Acinetobacter baumannii* | NZ_CP040048.1 | Complete C-I (CD Fused) |
| *Acinetobacter baumannii* | NZ_CP064203.1 | Complete C-I (CD Fused) |
| *Acinetobacter baumannii* | NZ_CP104448.1 | Complete C-I (CD Fused) |
| *Ralstonia solanacearum* | CP088234.1 | Complete C-I |
| *Ralstonia solanacearum* | NZ_CP115947.1 | Complete C-I |
| *Klebsiella pneumoniae* | NZ_CP159675.1 | Complete C-I (CD Fused) |
| *Klebsiella pneumoniae* | NZ_CP129740.1 | Complete C-I (CD Fused) |
| *Klebsiella pneumoniae* | NZ_CP129873.1 | Complete C-I (CD Fused) |
| *Burkholderia vietnamiensis* | JAKFAE010000004.1 | Complete C-I |
| *Mycobacterium intracellulare* | NZ_CP012886.2 | Complete C-I |
| *Citrobacter freundii* | CP048385.1 | Complete C-I (CD Fused) |
| *Klebsiella aerogenes* | LR134127.1 | Complete C-I (CD Fused) |
| *Komagataeibacter saccharivorans* | NZ_CP023037.1 | Complete C-I (CD Fused) |
| *Komagataeibacter saccharivorans* | NZ_CP036405.1 | Complete C-I (CD Fused) |
| *Tsukamurella tyrosinosolvens* | LR134465.1 | Complete C-I |
| *Legionella adelaidensis* | LR134433.1 | Complete C-I |
| *Paenibacillus cellulosilyticus* | CP054613.1 | C-I like (EF/EFG missing) |
| *Salmonella enterica* | NZ_CP087508.1 | Complete C-I (CD Fused) |
| *Salmonella enterica* | NZ_CP087512.1 | Complete C-I (CD Fused) |
| *Salmonella enterica* | NZ_CP087553.1 | Complete C-I (CD Fused) |
| *Sinorhizobium meliloti* | CP090106.1 | Complete C-I |
| *Sinorhizobium* sp. C101 | NZ_CP104135.1 | Complete C-I |
| *Sinorhizobium* sp. M103 | NZ_CP104127.1 | Complete C-I |
| *Sinorhizobium sp. K101* | NZ_CP104131.1 | Complete C-I |
